# The one-week automated genome-wide optical pooled screen

**DOI:** 10.64898/2026.04.15.718742

**Authors:** Bryce Kirby, Matteo Di Bernardo, Iain M. Cheeseman, Paul C. Blainey

## Abstract

Optical pooled screens (OPS) are bottlenecked by labor-intensive *in situ* sequencing and analysis protocols. Here we present OttoSeq, an automated OPS platform combining the Otto2 fluid handling system with the Brieflow analysis pipeline. We utilized OttoSeq to complete a genome-wide cell painting screen in eight days, sampling more than 5 million high-quality cells across 21,732 gene knockout perturbations (224 cells per gene) and interpreting 320 functional gene clusters.

## Introduction

Optical pooled screening (OPS) enables microscopy-based high-dimensional, single-cell phenotyping of CRISPR perturbation libraries. By reading out genetic barcodes through *in situ* sequencing by synthesis directly in fixed cells, OPS captures spatial and morphological information with high fidelity^1–3^ and offers a scalable alternative to next-generation sequencing (NGS)-based approaches such as Perturb-seq^4,5^. However, two bottlenecks have restricted OPS to specialist laboratories. First, *in situ* sequencing requires 1–2 hours of manual fluid handling per plate-base cycle, demanding weeks of expert hands-on labor. Second, the computational scripts most commonly used to process OPS data generalize to only a limited extent and require project-specific customization by experts^1–3,6^. While an integrated analysis solution has been recently reported^7^, it has not yet been benchmarked for rapid turnaround.

Production and primary readout of an optical pooled screen at genome-wide scale has thus required many months of hands-on labor and analysis. Here we present OttoSeq, an integrated system that addresses both bottlenecks by pairing the Otto2 on-microscope automated sequencing system with Brieflow^7^, a modular open-source analysis pipeline (Fig. 1a). To benchmark OPS throughput with OttoSeq, we completed a genome-wide cell painting screen in only eight days during which we generated cell painting data (20× magnification), amplified barcodes, and performed both the automated Otto2 *in situ* sequencing readout and automated Brieflow processing including MozzareLLM AI interpretation (Fig. 1b).

**Figure 1.**
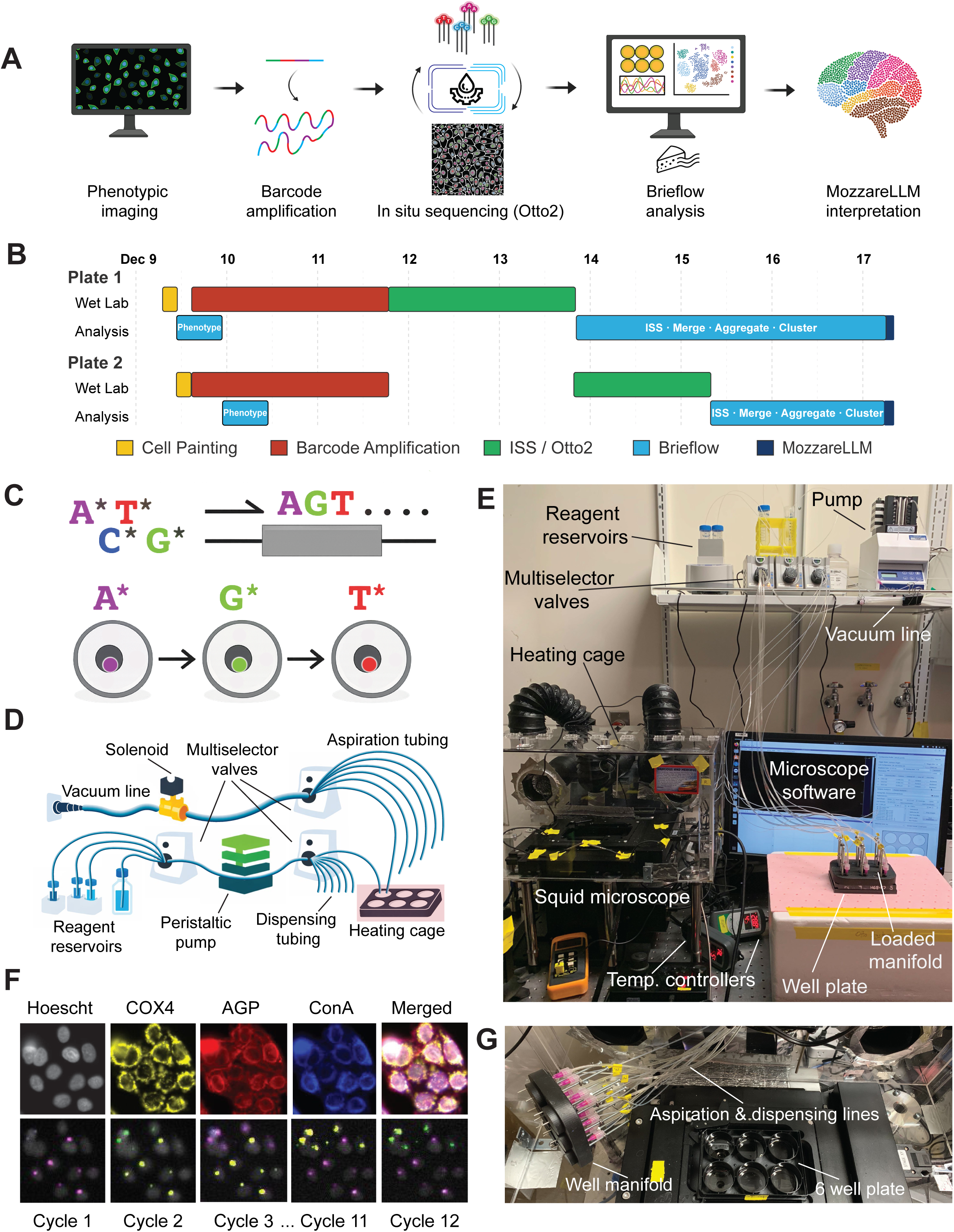
Otto2: an automated on-microscope in situ sequencing system. a, Schematic summary of the OttoSeq workflow. Cells are phenotypically imaged, perturbation barcodes are amplified, and plates undergo automated in situ sequencing on the Otto2 system. Images are processed through Brieflow for feature extraction, barcode calling, and clustering, and clusters are interpreted via MozzareLLM. Figure made with Biorender. b, To-scale timeline of the eight-day genome-wide screen by phase and plate, beginning with fixation and cell painting at 80 hours post-Cas9 induction. c, Schematic of the *in situ* sequencing-by-synthesis process over three cycles. The four nucleotide species are supplied as reversibly terminated, fluorophore-labeled nucleotides, with each base carrying a distinct fluorophore. Only a single nucleotide can be incorporated at each extension site per cycle. After nucleotide incorporation, the sample is imaged to identify the incorporated base. A cleavage step then removes the fluorophore and terminating group, regenerating the 3′ end and permitting incorporation of the next base in the following cycle. d, Schematic of the Otto2 fluid handling system. Reagents flow from cooled (4°C) or room-temperature reservoirs through multiselector valves and an inline peristaltic pump to individual wells in the well manifold and plate on the heated stage (45°C), then are removed from the plate through the vacuum line via a well multiselector valve when a solenoid valve is opened. e, Photograph of the Otto2 system preparing for calibration, with labeled parts. f, Example 20× crops of four-channel and merged cell painting images and 4× crops of five-channel merged SBS images across cycles, demonstrating relevant phenotypic stains and consistent, usable spot quality. g, Photograph of a plate ready to accept the well manifold with dispensing and aspiration lines secured for sequencing.

## Results

To address the challenges posed by existing protocols calling for 10+ cycles of labor-intensive *in situ* sequencing (Fig. 1c), we constructed Otto2, a minimal-cost, high-fidelity positive displacement pump system designed for rapid automated *in situ* sequencing in the six-well plate format (Fig. 1d,e). The system was designed to replicate manual *in situ* sequencing operations within a timeline comparable to human-operated benchmarks and produce base qualities sufficient for demultiplexing large scale screens. Otto2 performs semi-parallelized dispensing and aspiration operations lasting seconds, arranged in optimized step series addressed to individual wells. A single automated *in situ* cycle replaces 1 hour and 30 minutes of manual operations specified in previously reported protocols^3,6^. Otto2 is an on-microscope sequencer: tubing from the system hardware is threaded into a custom heating box integrated with the Cephla Squid+ fluorescence microscope^8^, holding plates at 45°C for chemistry operations excepting a final post-incorporation wash and addition of imaging buffer, which occur in each cycle at room temperature. To operate OttoSeq, the user loads a 3D-printed manifold into the wells of a six-well plate of cultured cells, positions it on the microscope, and secures dispensing and aspiration lines (Fig. 1g). Thus loaded, the plate is secure for an arbitrary number of sequencing and imaging cycles. By eliminating per-cycle manual fluid handling and by processing cycles on-scope, Otto2 reduces hands-on labor from ∼90 minutes to ∼10 minutes per cycle, enabling near-unattended and continuous operation.

Each plate was imaged at 20× magnification in four spectral channels (Fig. 1f) for cell painting—Hoechst (DNA), COX4 (mitochondria), AGP (actin and Golgi), and Concanavalin A (endoplasmic reticulum)—an OPS-optimized variation of the original cell painting panel^9^.

Following cell painting and barcode amplification (Methods), each of the two screening plates were loaded for 12 cycles of *in situ* sequencing using Otto2 at 4× magnification (Fig. 1f). During each cycle, following 10 minutes of setup and quality control, the Otto2 system ran autonomously for ∼2.5 hours including imaging.

Matching automated data generation speed required an adaptable analysis pipeline. Brieflow, a modular open-source pipeline for OPS data, processed raw images^7^. Brieflow’s modular architecture was essential to achieving the 8-day overall turnaround: because individual processing steps—segmentation, feature extraction, barcode calling, spatial merging, clustering—are implemented as independent, swappable modules, improvements could be deployed without modifying upstream or downstream steps. The genome-wide screen required on-the-fly adaptations to two modules during the screen: enhanced SBS cycle alignment to accommodate unexpectedly large inter-cycle plate position offsets, and a filtered analysis arm designed to efficiently subcluster on only bootstrap-significant perturbations given the scale of the library (Methods). These updates allowed the analysis pipeline to keep pace with automated hardware. Bug fixes and performance improvements (Supplementary Table 1) were merged back into the main codebase. Brieflow processed 13,210,438 barcoded cells and 15,647,674 phenotyped cells across two six-well plates, extracting 1,670 morphological features from each phenotyped cell. After spatial merging, deduplication, and quality filtering, 5,198,240 mapped cells were retained across 21,732 perturbations with a median of 224 cells per gene (Fig. 2a–c, Supplementary Table 2). These coverage statistics confirm that OttoSeq preserved statistical power for genome-wide morphological profiling under the compressed eight-day timeline.

**Figure 2.**
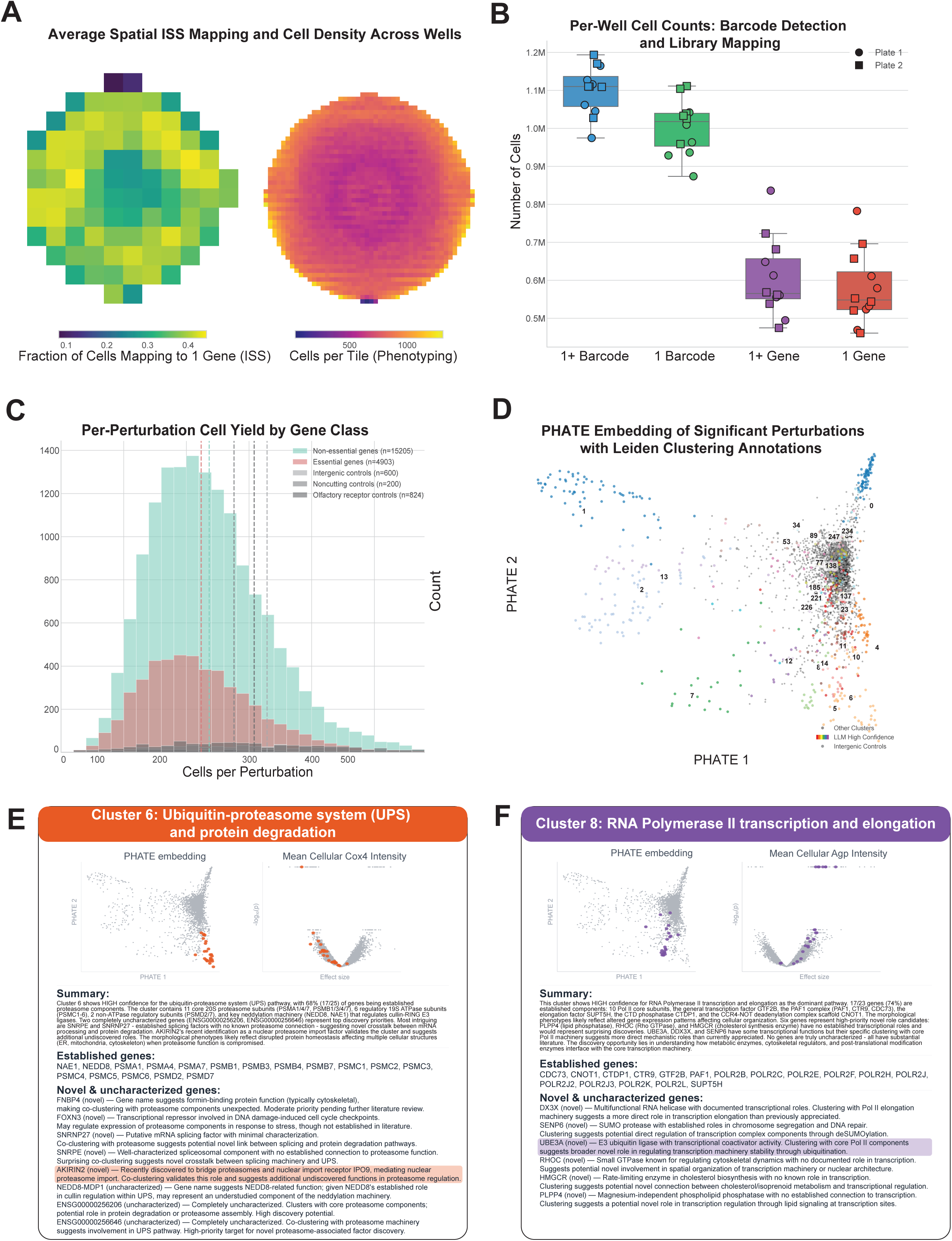
Automated analysis and genome-wide screen results. a, Spatial quality control averaged across all wells, showing the fraction of cells mapping to exactly one gene and cell density per tile. b, Counts per well for per-cell mapping categories. On average, 73.1% of cells mapped to at least one barcode, 66.4% to exactly one barcode, 40.2% to at least one gene, and 38.3% to exactly one gene. c, Distribution of cells per perturbation stratified by gene class, with median thresholds indicated by vertical dotted lines. d, PHATE embedding of 3,715 bootstrap-significant perturbations, with MozzareLLM high-confidence clusters labeled by Leiden cluster number. e-f, MozzareLLM interpretation cards for other core cellular processes: Cluster 6 (ubiquitin-proteasome system, 68% established) and Cluster 8 (RNA Polymerase II transcription and elongation, 74% established), showing PHATE localization, representative volcano plot, and default LLM interpretation output.

A final challenge was biological interpretation, which typically requires weeks to months of manual literature curation. We applied MozzareLLM^7^, an LLM-driven tool that generates biological summaries from perturbation cluster data within hours. Bootstrap significance testing identified 3,715 perturbations with significant morphological effects, which were organized into 320 clusters (Fig. 2d, Supplementary Table 3). MozzareLLM then assessed each cluster for biological coherence, assigning pathway confidence tiers—27 high, 168 medium, and 125 low—reflecting the strength of evidence that a cluster captured a unified biological process. The high-confidence clusters (610 genes, 75% essential in HeLa) recover well-characterized cellular machinery - biologically validating accurate clustering of functionally related genes - alongside a much smaller number of genes (152) in unexpected functional contexts which represent putative discoveries (Supplementary Table 4). Clusters mapping to ubiquitin-proteasome degradation (Cluster 6, Fig. 2e) and RNA Pol II transcription (Cluster 8, Fig 2f) recover recently validated genes such as AKIRIN2, mediating nuclear proteasome import^10^, and novel findings such as UBE3A, an Angelman syndrome-associated E3 ligase clustering with core Pol II machinery. A select few genes in these clusters were not previously assayed by HeLa essential gene CRISPR libraries^3^, making them promising subjects for further investigation (Supplementary Tables 3, 4).

## Discussion

Although OPS offers high-content screening of genome-wide CRISPR libraries, it has previously been bottlenecked by library demultiplexing and analysis, with screens routinely requiring many weeks of data generation and months of analysis. Our screen required only eight days with two researchers and consisted largely of hands-off automated runtime, drastically reducing cost and user burden. Several considerations remain: on-scope sequencing presents difficulties unique to fluidic-imaging integration—including plate position offsets from tubing drag, material compatibility with epifluorescence imaging, and parallelization limited to one plate per microscope—and per-cycle Otto2 runtime is ∼50% longer than manual cycling. Automated *in situ* sequencing with OttoSeq unlocks previously unattainable throughput for high-content perturbation screening and suggests the possibility of routine future access to high-speed, low-cost perturbation screening for many biomedical researchers.

## Methods

### Library and transduction

The genome-wide CROPseq-multi CRISPR-KO library targeted 20,553 genes (union of GENCODE, RefSeq, and CHESS annotations, including 412 olfactory receptors) with 41,906 dual-guide constructs in the CROPseq-multi-puro-v2 vector^11^. Each gene was targeted by two dual-guide constructs carrying four unique CRISPick-selected guides paired by pick-sum ordering (guides 1+4 and 2+3) with random position assignment^12–14^. Controls comprised 600 intergenic-targeting constructs (paired guides targeting the same chromosome, 100–10,000 bp apart, with comparable on-target efficacy and specificity scores to gene-targeting guides), 200 nontargeting constructs, and 824 constructs targeting 412 olfactory receptor genes. Each construct carried a pair of iBARs with a guaranteed Levenshtein edit distance of 3 across 12 sequencing cycles, enabling error-corrected barcode calling; observation of either iBAR alone is sufficient to identify a construct. Lentivirus was produced by transfecting HEK-293T cells seeded at 15 × 10⁶ cells in a T175 cm² flask in DMEM supplemented with 10% FBS and 1% penicillin-streptomycin 30 hours pre-transfection. Media was changed to 20 mL virus production medium (IMDM supplemented with 20% heat-inactivated FBS) 7–8 hours pre-transfection. Cells were transfected with 20 µg plasmid library, 3.62 µg pCMV-VSV-G, 8.28 µg psPAX2, and 95.8 µL FuGENE 4K in Opti-MEM to a final volume of 2 mL. Media was changed to 55 mL fresh virus production medium 16–17 hours post-transfection. Viral supernatant was harvested at 48 hours post-transfection, 25 mL fresh medium was added, and a second harvest was collected at 72 hours. Supernatant was 0.45 µm filtered and stored at −80°C. For the screen, HeLa-TetR-Cas9 (A7) cells were transduced with the sgRNA library in CROPseq-multi-v2-Puro and selected with 2 μg/mL puromycin (Sigma Aldrich P45-12) for 4 days.

### Cell culture

Media comprised DMEM GlutaMAX (Gibco 10569010) with 10% (v/v) heat-inactivated fetal bovine serum (MilliporeSigma F4135), 100 U/mL Penicillin, and 100 μg/mL Streptomycin (Gibco 15140122). Cells were thawed, added dropwise to warmed media, spun down at 400g x 5 minutes, resuspended, and seeded at half-confluency into T225 cm² culture flasks (Fisher Scientific 159934) at scale sufficient for 1000x gene-perturbation representation. Following a 48-hour incubation, an exchange for media containing 2ug/mL doxycycline was performed to induce Cas9 activity. Following a 24-hour incubation, cells were washed once with 1X PBS, passaged with TrypLE (Thermo Fisher Scientific 12604013), and seeded in two six-well plates (P06-1.5H-N) at a density of 275K cells per well in 2mL of media containing 2ug/mL doxycycline. Following a 56-hour incubation (for 80 hours total of doxycycline exposure), plates were washed once with 1X PBS, fixed in 4% formaldehyde (Electron Microscopy Sciences 15714) in 1X PBS (Ambion AM9625) for 30 min at room temperature, and washed three times with 1X PBS.

### Cell painting

Plates were incubated with cell painting mix containing 1:5000 Hoechst (Thermo Fisher Scientific 62249), 1:250 COX4 (ProteinTech CL488-11242), 1:1000 Phalloidin (Thermo Fisher Scientific A34055), 1:7500 WGA (Thermo Fisher Scientific W32464), and 1:500 Concanavalin A (Thermo Fisher Scientific C21421) in 2.5% BSA (LCG 1900-0016) and 0.5% Tween-20 (VWR 100216-360) in 1X PBS (Ambion AM9625) for 45 minutes at room temperature and washed three times with 0.05% Tween-20 in 1X PBS. Imaging setup consisted of a Cephla Squid+ microscope with a 5 line laser engine (405 nm, 470 nm, 550 nm, 640 nm, 730 nm), a motorized filter wheel, 850 nm laser autofocus and a camera with Sony IMX571 sensor (3.76 um pixel size, 4168 x 4168 square FOV). Excitation filter was Chroma ZET405/470/555/640/730 and dichroic was Semrock FF421/491/567/659/776-Di01. Plates were imaged in 4 channels per field of view (Hoechst: 405nm excitation with Semrock FF01-441/511/593/684/817 emission filter, henceforth multiband emission filter, COX4: 470nm excitation with Chroma ET525/50m emission filter, AGP: 550nm excitation with multiband emission filter, Concanavalin A: 640nm excitation with multiband emission filter) using a 20× objective (Olympus Plan Apo 20×/0.8) with no z-stacking at 2×2 binning.

### T7-based barcode amplification

Barcode amplification utilizing the T7-IVT (PerturbView) in situ amplification protocol with the Crop-Seq Multi vector is described in detail elsewhere^11,15^. Briefly, barcode transcripts were generated and converted to localized rolling-circle amplicons as follows. Cells were fixed in 4% (v/v) formaldehyde (Electron Microscopy Sciences 15714) in 1X PBS (Ambion AM9625) for 30 minutes at room temperature and washed twice with PBS. Cells were permeabilized in 70% ethanol for 30 minutes, washed three times with PBS-T (1X PBS + 0.05% Tween-20; VWR 100216-360), and decrosslinked in 0.1 M sodium bicarbonate and 0.3 M NaCl at 65 °C for 4 hours followed by three PBS-T washes. All reactions were performed in heat-compatible imaging plates (Cellviz P96-1.5H-N). In-situ transcription was performed in T7 reaction buffer (NEB E2040L) containing 10 mM each NTP (NEB E2040L), 5 mM DTT (NEB E2040L), 0.4 U/µL RiboLock RNase inhibitor (Thermo Fisher Scientific EO0384), and T7 RNA polymerase mix (NEB E2040L - M0255AVIAL) diluted to 0.2X relative to the manufacturer’s recommendation. Reactions were incubated at 37 °C for 12 hours. Samples were washed three times with PBS-T and post-fixed in 3% formaldehyde (Electron Microscopy Sciences 15714) and 0.1% glutaraldehyde (Electron Microscopy Sciences 16120) for 30 minutes, followed by quenching for 1 minute in pH 8 0.2 M Tris-HCl (Thermo Fisher Scientific AM9855G) and three PBS-T washes. Reverse transcription was carried out in 1X RevertAid RT buffer (Thermo Fisher Scientific EP0452) containing 250 µM dNTPs (NEB N0447L), 1 µM biotinylated reverse-transcription primer (Integrated DNA Technologies), 200 µg/mL recombinant albumin (NEB B9200S), 0.8 U/µL RiboLock RNase inhibitor (Thermo Fisher Scientific EO0384), and 4.8 U/µL RevertAid H-minus Reverse Transcriptase (Thermo Fisher Scientific EP0452). Reactions were incubated at 37 °C for 12 hours. Samples were washed twice with PBS-T and incubated for 15 minutes with 20 µg/mL streptavidin (NEB N7021S) and 100 µg/mL recombinant albumin (NEB B9200S) in PBS, followed by two PBS-T washes and postfixation in 3% formaldehyde (Electron Microscopy Sciences 15714) and 0.1% glutaraldehyde (Electron Microscopy Sciences 16120) for 30 minutes. Padlock probe gap-fill and ligation were performed in 1X Ampligase buffer (Lucigen A3210K) containing 50 nM dNTPs (NEB N0447L), 0.1 µM each padlock probe (Integrated DNA Technologies), 200 µg/mL recombinant albumin (NEB B9200S), 0.4 U/µL RNase H (Enzymatics Y9220L), 0.02 U/µL TaqIT polymerase (Enzymatics P7620L), and 0.5 U/µL Ampligase (Lucigen A3210K). Samples were incubated in gap-fill and ligation solution at 37 °C for 5 minutes followed by 45 °C for 90 minutes, then washed twice with PBS-T. Rolling circle amplification (RCA) was performed in 1X Phi29 buffer (Thermo Fisher Scientific EP0091) containing 250 µM dNTPs (NEB N0447L), 5% glycerol (MilliporeSigma G5516), 200 µg/mL recombinant albumin (NEB B9200S), and 1 U/µL Phi29 DNA polymerase (Thermo Fisher Scientific EP0091) for 12 hours at 30 °C. Samples were washed twice with PBS-T, then 1 µM sequencing primers (Integrated DNA Technologies) were hybridized in 2X SSC (Ambion AM9763) with 10% (v/v) formamide (Thermo Fisher Scientific 4650) for 30 minutes at room temperature and washed twice with PBS-T prior to *in situ* sequencing.

### Otto2 system assembly and operation

The Otto2 system was constructed from off-the-shelf parts. The Otto2 system comprised 3 multi selector valves (MXX778-605) and 1 standard flow eight-channel head peristaltic pump (Gilson F155006) wired for programmatic control to an Arduino Mega 2650 microcontroller. Cooling blocks (Fisher Scientific 14-955-220) included aluminum 50mL conical inserts. Tubing was chemical-resistant 1/16” OD tubing (McMaster-Carr 5583K51) for reagent and dispensing tubing, ⅛” OD tubing (McMaster-Carr 9117T8) for aspiration tubing, and 1.02mm ID PVC tubing for pump tubing (Gilson F117938). Leur locks were used to connect 16-gauge blunt-end sample needles (McMaster-Carr 75165A753) to 1/16” (McMaster-Carr P-836) and ⅛” (McMaster-Carr P-831) tubing. The Otto2 cage was constructed from cast acrylic sheets (McMaster-Carr) with ¼” sheets used for walls and ⅛” sheets used for floor and wall, cut to size with a laser cutter (Epilog Fusion Pro) and sealed using industrial acrylic cement followed by silicone sealant. The cage was temperature controlled using two parallel heating systems (right/left), each consisting of a 600 watt PTC fan heater placed within a closed ducting system composed of 3” supply and return ducts and upstream of a dedicated heating controller (Inkbird ITC-308). A 3/8’ 12V Solenoid Valve (U.S. Solid), controlled via an optocoupler relay switch, was laid inline into ⅜” heavy-wall vacuum tubing downstream of a 1L vacuum trap sealed with a ⅜” drilled rubber bung. Vacuum tubing was connected to an in-house vacuum wall spigot and adapted on the upstream end to match ⅛” OD aspiration multivalve selector input. The well manifold was constructed using a Formlabs 3 SLA resin printer with Black v4.1 resin (RS-F2-GPBK-41) and cured according to manufacturer instructions. #6-32 flat-headed screws threaded with flange nuts, used to secure tubing needles, were screwed down into matching holes in the manifold surface until resistance was met. *In situ* cycling operations with the Otto2 system proceeds as follows: a) multi-selector reagent valve is set to the appropriate reagent (incorporation mix, cleavage mix, incorporation wash buffer, or imaging buffer); b) reagent is drawn up from chemistry reservoirs through a dynamically pre-set reagent multiselector valve and inline peristaltic pump via positive displacement action through a dynamically pre-set dispensing multiselector valve and into a selected plate well via dispensing tubing and needle; c) an aspiration multiselector valve is set to a specified well; d) the solenoid valve is opened and well-specific aspiration occurs through the aspiration needle and tubing. Operations occur in a semi-parallel fashion such that while well B undergoing aspiration, well A is undergoing dispensing, followed by well C undergoing aspiration while well B is undergoing dispensing. Once dispensed, sequencing reagents were followed by an air bubble and then wash reagent to keep reagents unmixed. Otto2 control software was written in Arduino and accessed via publicly available Arduino IDE.

### Automated in situ sequencing protocol

The first incorporation cycle was conducted manually to decouple barcode prep from automation functionality in a fashion similar to previous reports^2^ but utilizing diluted incorporation mix to further reduce screen cost and offset the higher minimum reagent volumes necessary for automated function compared with the manual protocol. Briefly, incorporation mix (reagent 1) was extracted from thawed Illumina MiSeq Nano v2 kits (Illumina MS-103-1003); wells were incubated with incorporation reagent diluted 2:1 in 1X ThermoPol buffer (Thermo Fisher Scientific 62249) for 7.5 minutes on a flat-top thermal cycler at 45C, then exchanged three times in PR2 buffer (Illumina MS-103-1003), then washed quickly three times with PR2, then incubated for 5 minutes at 45C on a flat-top thermal cycler five times, then washed once in PR2, then incubated for 5 minutes in 2X SSC (Thermo Fisher Scientific AM9763) with 1:2000 Hoechst (Thermo Fisher Scientific 62249). Otto2 system was calibrated by setting correct pump time for 1mL dispensation and by validating successful dispensing, aspiration, and reagent addition operations. Once the heating box reached equilibrium at 45C (15 minutes) and cooling blocks were cooled to 4C, the plate was loaded with the manifold and positioned on the microscope stage. Needle ends of dispensing and aspiration tubing were loaded and secured into correct positions in manifold with one dispensation and aspiration line per well. Per base cycle, 7.5mL of 1.5X diluted incorporation mix (prepared as previously described), 7.5mL of pure cleavage mix, and 50mL of 2X SSC with 1:2000 Hoechst were placed in separate 50ml conicals in cold block inserts with respective 1/16” OD reagent tubing secured into to the bottom of each conical. A bottle containing 300mL of PR2 buffer was secured with 1/16” OD reagent tubing in the same fashion. Cycle operation was launched via serial connection with the microcontroller unit. Cycle operation was hands-off; the user observed two diagnostic dispensing and aspiration operations across wells at cycle onset and observed imaging collection to validate correct performance. Cycle operation sequence for each sequencing reagent following two diagnostic washes (1mL/well) proceeded as follows, first for cleavage then incorporation: reagent addition (0.98mL); 1 exchange wash (1mL dilution pre-1mL wash); 1 quick wash (0.75mL); 1 increased volume wash (3 mL); 4 standard washes (1mL). Reagent addition incubations were 7.5 minutes, cleavage wash incubations were 1 minute, and incorporation wash incubations were 5 minutes. All washes were conducted with PR2. The cycle concluded with a 12 minute incubation, a 2X addition (1mL) of imaging buffer and a final addition of imaging buffer (3 mL). Otto2 system held temperature at 45C for all operations through final 5 minute wash and at room temperature for final 12 min incubation, imaging buffer operations, and imaging. Conclusion of an Otto2 sequencing cycle automatically triggered onset of imaging via custom integration in Squid source software. Images were collected on the Cephla Squid+ using the aforementioned imaging setup in 5 channels per field of view (Hoechst: 405nm excitation with multiband emission filter, G: 550nm excitation with FF01-578/21 emission filter, T: 550nm excitation with FF01-624/40 emission filter, A: 640nm excitation with FF01-680/42 emission filter, C: 640nm excitation with FF01-732/68 emission filter) using a 4× Olympus objective (Olympus Plan Apo 4×/0.16) at 1×1 binning.

### Data transfer and compute

Raw images were transferred from the microscope workstation to Google Cloud Storage using parallelized gsutil rsync (12 processes × 8 threads). Phenotype images uploaded in approximately 55 minutes per plate (∼1 hour 50 minutes total); SBS cycle images uploaded in approximately 2 hours 48 minutes per plate (∼5 hours 36 minutes total across both plates).

Images were downloaded from GCS to the compute instance in approximately 10 minutes for phenotype data and 13 minutes per SBS cycle, benefiting from GCS–GCP colocation. On GCP, Brieflow preprocessing completed in approximately 25 minutes per plate, phenotype segmentation and feature extraction required approximately 3 hours per plate (∼6 hours 11 minutes total), SBS processing required approximately 3 hours per plate (plate 1 required additional time due to rotation alignment debugging), spatial merging completed in approximately 18 minutes, aggregation (including bootstrapping) in approximately 4 hours 19 minutes, and clustering across multiple resolutions in approximately 1 hour 55 minutes.

MozzareLLM interpretation of 320 clusters completed in approximately 1 hour 41 minutes. Thus, total runtime was approximately 31 hours, although this does not include parameter setting, which is done in user facing notebooks (which takes <30 minutes per module). All computation was performed on a GCP c4d-standard-384 instance (384 vCPUs, 1,488 GB RAM). Complete stage-by-stage runtimes are provided in Supplementary Table 5 and a detailed analysis timeline in Supplementary Table 6.

### Brieflow analysis pipeline

Brieflow (v1.4.6)^7^ was used for all computational analysis. The genome-wide screen served as a stress test for the pipeline, and the resulting changes fell into two categories. First, general bug fixes and performance improvements — including parallelized SBS parameter search, corrected IC field image combination, multi-mode evaluation support, and a windowed alignment parameter — were merged back into Brieflow’s main codebase (PR #177, 10 commits; Supplementary Table 2), directly benefiting future users. Second, screen-specific adaptations — rotation-aware SBS alignment and a parallel filtered clustering arm — were developed on a dedicated branch (Supplementary Table 7) and are described below.

### SBS cycle alignment

Automated on-scope fluid handling introduced inter-cycle drift exceeding what standard translation-only registration could correct. To address this, we added a rotation-and-translation alignment method to Brieflow’s SBS alignment module. For each cycle, the DAPI channel was used as a reference. A rough translational offset was first computed on a windowed center region via phase cross-correlation, then applied to the full image. A Laplacian of Gaussian filter (sigma = 3.0) was applied to enhance spot features, and rotation was estimated by phase cross-correlation in log-polar space. The rotation was applied to the original source image before a final translational offset was computed in the rotated coordinate frame. Rotations exceeding 5° were clamped to zero as likely spurious. For tiles where automatic registration failed due to large drift, manual coarse offsets could be specified per cycle and applied as a pre-alignment step before the automatic fine alignment. An alternative edge-based alignment using Canny edge detection (sigma = 2.0) on DAPI was also implemented for robustness to signal degradation over cycles. All alignment methods were added to the shared alignment module without modification to downstream barcode calling or feature extraction. We sequenced deeply enough to enable error correction mode, a feature of the sparse sampling in Levenshtein space which allows barcode matching within a distance of two bases.

### Feature extraction, merging, and aggregation

Segmentation, feature extraction (1,670 CellProfiler-derived features^16^ per cell), spatial merging, deduplication, and aggregation were performed using Brieflow’s standard modules as previously described^7^. Bootstrap significance testing used 10,000 simulations per gene against non-targeting controls, with a z-score threshold of 0.3 and FDR threshold of 0.5, yielding 3,715 significant perturbations from 21,732 total (Supplementary Table 4).

### Filtered clustering

To focus downstream analysis on biologically meaningful perturbations, bootstrap-significant genes were used to filter the aggregated feature table. PHATE embedding^17^ and Leiden clustering^18^ were run at multiple resolutions (10.0–15.0) and benchmarked against CORUM^19^ and KEGG^20^ annotations. Resolution 10.0 was selected, yielding 320 clusters with 15 CORUM-enriched and 8 KEGG-enriched clusters (Supplementary Table 3).

### MozzareLLM interpretation

Cluster gene lists and associated UniProt functional annotations were passed to MozzareLLM^7^. The screen context prompt was varied slightly to inform the LLM API calls of the markers used in the screen. MozzareLLM queried Claude Sonnet 4.5 (claude-sonnet-4-5-20250929) to generate structured biological interpretations for each of 320 clusters, including dominant biological processes, pathway assignments, and per-gene classifications (established function, novel role, or uncharacterized). Confidence was assigned as high, medium, or low based on the coherence of gene functions within each cluster (Supplementary Tables 3, 4).

## Availability of data and materials

Brieflow is available at https://github.com/cheeseman-lab/brieflow under an open-source license. The analysis configuration and scripts used for this screen, including scripts for figure generation, are available at https://github.com/cheeseman-lab/whitney-analysis. MozzareLLM is available at https://github.com/cheeseman-lab/mozzarellm. The raw data supporting the conclusions of this article is available at the BioImage Archive: https://www.ebi.ac.uk/biostudies/bioimages/studies/S-BIAD3096. We also provide an interactive viewer for exploring the data and all quality control metrics at https://whitney.broadinstitute.org/.

## Authors’ contributions

BK conceived of the work and developed the current Otto2 fluid handling system. MD developed the Brieflow computational pipeline and MozzareLLM framework. BK and MD executed the screen, performed data analysis, and co-wrote the manuscript. IMC and PCB supervised the work and edited the manuscript.

## Supporting information

Well insert print file

## Acknowledgements

We thank Anna Le, Owen Andrews, Sami Farhi, and Tridib Biswas for their foundational contributions to developing and testing the automation platform. We thank Russell Walton for piloting, designing, and generating the perturbation library. We thank Heather Keys, Kuan-Chung Su, Kaitlyn Manzer, and Russell Walton for lentivirus production, transduction, and cell line development. We especially thank Cephla for loaning the Squid+ microscope, as well as Hongquan Li and You Yan for their continuous support and assistance in configuring the imaging platform. The HeLa cell line was used in this research. Henrietta Lacks, and the HeLa cell line that was established from her tumor cells without her knowledge or consent in 1951, have made significant contributions to scientific progress and advances in human health. We are grateful to Lacks, now deceased, and to the Lacks family for their contributions to biomedical research.

## References

1. Feldman, D. et al. Optical Pooled Screens in Human Cells. Cell 179, 787–799.e17 (2019).

2. Feldman, D. et al. Pooled genetic perturbation screens with image-based phenotypes. Nat Protoc 17, 476–512 (2022).

3. Funk, L. et al. The phenotypic landscape of essential human genes. Cell 185, 4634–4653.e22 (2022).

4. Dixit, A. et al. Perturb-Seq: Dissecting Molecular Circuits with Scalable Single-Cell RNA Profiling of Pooled Genetic Screens. Cell 167, 1853–1866.e17 (2016).

5. Replogle, J. M. et al. Mapping information-rich genotype-phenotype landscapes with genome-scale Perturb-seq. Cell 185, 2559–2575.e28 (2022).

6. Carlson, R. J., Leiken, M. D., Guna, A., Hacohen, N. & Blainey, P. C. A genome-wide optical pooled screen reveals regulators of cellular antiviral responses. Proceedings of the National Academy of Sciences 120, e2210623120 (2023).

7. Di Bernardo, M. et al. Brieflow: An Integrated Computational Pipeline for High-Throughput Analysis of Optical Pooled Screening Data. Preprint at 10.1101/2025.05.26.656231 (2025).

8. Li, H. et al. Squid: Simplifying Quantitative Imaging Platform Development and Deployment. Preprint at 10.1101/2020.12.28.424613 (2020).

9. Cimini, B. A. et al. Optimizing the Cell Painting assay for image-based profiling. Nat Protoc 18, 1981–2013 (2023).

10. de Almeida, M. et al. AKIRIN2 controls the nuclear import of proteasomes in vertebrates. Nature 599, 491–496 (2021).

11. Walton, R. T. et al. CROPseq-multi: a universal solution for multiplexed perturbation in high-content pooled CRISPR screens. 2024.03.17.585235. Preprint at 10.1101/2024.03.17.585235 (2025).

12. Doench, J. G. et al. Optimized sgRNA design to maximize activity and minimize off-target effects of CRISPR-Cas9. Nature Biotechnology 34, 184–91 (2016).

13. Sanson, K. R. et al. Optimized libraries for CRISPR-Cas9 genetic screens with multiple modalities. Nature Communications 21, 5416 (2018).

14. Drepanos, L. M. et al. Balancing off-target and on-target considerations for optimized Cas9 CRISPR knockout library design. 2025.08.26.672375. Preprint at 10.1101/2025.08.26.672375 (2025).

15. Kudo, T. et al. Multiplexed, image-based pooled screens in primary cells and tissues with PerturbView. Nature Biotechnology 43, 1091–1100 (2025).

16. McQuin, C. et al. CellProfiler 3.0: Next-generation image processing for biology. PLoS Biol 16, e2005970 (2018).

17. Moon, K. R. et al. Visualizing Structure and Transitions in High-Dimensional Biological Data. Nature Biotechnology 37, 1482–92 (2019).

18. Traag, V. A., Waltman, L. & Eck, N. J. From Louvain to Leiden: Guaranteeing Well-Connected Communities. Scientific Reports 9, 5233 (2019).

19. Giurgiu, M. et al. CORUM: the comprehensive resource of mammalian protein complexes—2019. Nucleic Acids Research 47, D559–D563 (2019).

20. Kanehisa, M., Furumichi, M., Sato, Y., Matsuura, Y. & Ishiguro-Watanabe, M. KEGG: Biological Systems Database as a Model of the Real World. Nucleic Acids Research 53, 672–77 (2025).

